# Differentiation of Human induced Pluripotent Stem Cells to Authentic Macrophages using Fully Defined, Serum Free, Open Source Media

**DOI:** 10.1101/2020.10.27.357632

**Authors:** Alun Vaughan-Jackson, Szymon Stodolak, Kourosh H. Ebrahimi, Cathy Browne, Paul K. Reardon, Elisabete Pires, Javier Gilbert-Jaramillo, Sally A. Cowley, William S. James

**Author notes:** Correspondence: Alun Vaughan-Jackson @VaughanAlun William S. James.

## Abstract

**Summary:** Human iPSC and macrophages derived from them are increasingly popular tools for research into both infectious and degenerative diseases. However, as the field strives for greater modelling accuracy, it is becoming ever more challenging to justify the use of undefined and proprietary media for the culture of these cells. We describe here two fully defined, serum-free, open-source media for the culture of iPSC and differentiation of iPSC-derived macrophages. These media are equally capable of maintaining these cells compared to commercial alternatives. The macrophages differentiated in these defined media display improved terminally differentiated cell characteristics, reduced basal expression of induced anti-viral response genes, and improved polarisation capacity. We conclude that cells cultured in these media are an appropriate and malleable model for tissue resident macrophages, on which future differentiation techniques can be built.

## Introduction

Resident phagocytes are an evolutionarily conserved cell type in metazoans. In mammals, resident macrophages support tissue homeostasis through a wide range of specialized trophic, remodelling and defence functions, whose importance is illustrated by their failure in malignant, degenerative and infectious diseases (Gordon et al., 2014; Steinman and Moberg, 1994). In order to investigate the molecular pathways involved in the pathogenesis of both infectious diseases, such as those caused by HIV-1 (van Wilgenburg et al., 2013) and *Mycobacterium tuberculosis* (Härtlova et al., 2018), and degenerative diseases, especially Parkinson’s disease (Haenseler et al., 2017a; Lee et al., 2020) and Alzheimer’s disease (Brownjohn et al., 2018; Garcia-Reitboeck et al., 2018), we developed a pathophysiologically authentic, yet genetically tractable model of human tissue macrophages derived from pluripotent stem cells (Karlsson et al., 2008; van Wilgenburg et al., 2013). We have shown this model resembles a MYB-independent early wave of myelopoiesis *in vivo* that gives rise to resident macrophages in a number of tissues (Bian et al., 2020; Buchrieser et al., 2018). Our first method involved spontaneous differentiation of mesoderm from embryonic stem cells via embryoid bodies (EBs), followed by myeloid differentiation using M-CSF (CSF-1) and IL-3 in a serum-supplemented medium (Karlsson et al. 2008). Because this was not effective for all pluripotent stem cell lines, and in order to remove the undefined serum component, we subsequently developed a serum-free method, using BMP4, VEGF and SCF to promote mesodermal lineage and prime haemogenic endothelium differentiation during EB formation, and the serum-free X-VIVO™ 15 medium (Lonza) during the M-CSF/IL-3-directed myeloid differentiation stage (van Wilgenburg et al. 2013). While ours is not the only methodology for iPSC-derived macrophage differentiation, other methods require the use of serum, or proprietary media, or both. (Choi et al., 2009; Gupta et al., 2016; Kambal et al., 2011; Subramanian et al., 2011; Yanagimachi et al., 2013; Zhang et al., 2015; and reviewed by Lee et al., 2018)

With a greater focus than ever on ensuring research integrity in the life sciences through the adoption of open practices in data curation, publication and the sharing of materials (Cech et al., 2003; Morey et al., 2016; National Academy of Sciences, National Academy of Engineering (US) and Institute of Medicine (US) Committee on Science, 2009), it is desirable to replace proprietary media used for culturing and differentiating human pluripotent stem cells with ones that are both defined and fully open-source. The suppliers of both the iPSC culture medium Essential 8 (StemCell Technologies and Life Technologies) and X-VIVO™ 15 decline to publish the composition of these materials on commercial grounds. Consequently, the scientific community does not know whether components that might have a material effect on the physiology of the stem cells and macrophages being studied are present, nor their concentration. Examples could include, but are not limited to, anti-infectives, anti-inflammatories, cell-permeable iron-chelators and cryoprotectants. In macrophages, the balance between oxidative phosphorylation and aerobic glycolysis appears to drive a regulatory versus an inflammatory immune response (Kramer et al., 2014; Palmer et al., 2015). There is therefore a concern that, in the presence of excessively high concentrations of glucose and insulin, for example, cells might display abnormal rather than physiological responses to stimuli. Accordingly, we have sought to base replacement media on widely available and open-source materials, enabling others to reproduce our experiments conveniently, inexpensively, and without undue dependence on particular suppliers.

In this paper, we describe OXE8, a cost-effective and convenient, fully defined and open-source modification of E8 medium, based on well-defined commercial Advanced DMEM/F12, which performs at least as well as proprietary media. We also describe an alternative macrophage differentiation medium, OXM, which is both defined and open source. We show that OXM generates homogeneous macrophage precursors very comparable to those produced by our earlier methods, both phenotypically and transcriptionally. We note that these cells show signatures consistent with more complete terminal differentiation, including improved morphology and cell cycle arrest, and have lower basal levels of expression of interferon-inducible genes while being more responsive to inflammatory stimulation. This method therefore generates cells that are even closer models of the “surveilling” state of homeostatic tissue macrophages.

## Results

### Culture of iPSC in fully defined, open source medium, OXE8

To culture iPSC-derived macrophages using fully defined, open-source media, we first defined the medium for culturing of iPSC. We devised our own fully defined, open-source, easy to make and relatively cheap alternative (approximately 1/5 the cost) to E8 medium for culturing iPSC, based on Advanced DMEM/F-12 (aDMEM/F-12) (Gibco), which we named OXE8. All components of aDMEM/F-12 and their concentrations can be found at the supplier’s site. Of the 8 components described by Chen et al. 2011 (Chen et al., 2011), insulin (1.72 μM), selenium (Sodium selenite, 29 nM), L-ascorbic acid (L-ascorbic acid-2-phosphate magnesium, 86 μM), and transferrin (7.5 mg/L) are already components of aDMEM/F-12. FGF-2, and TGFβ, and additional L-ascorbic acid were supplemented independently. In addition, Heparin was added to stabilise the FGF-2 (Chen et al., 2012; Furue et al., 2008), and the medium was supplemented with human serum albumin (HSA). The medium was buffered with HEPES. For the full composition, see Supplementary Table 1.

To evaluate the effect of culturing iPSC in OXE8, cells were cultured over five passages (18 days), passaging when cells were approximately 80-90% confluent. Growth and phenotype of iPSC cultured in OXE8 were compared to the current commercial culture media mTeSR™ (StemCell Technologies) and Essential 8 (Gibco), herein E8. Cell morphology, number, and viability were not significantly different between culture conditions across the five passages (Figure 1A-D). Expression of pluripotent stem cell markers NANOG and TRA-1-60, was the same across all 3 culturing conditions, as assessed by flow cytometry (Figure 1E-F). Together, this analysis demonstrates that OXE8 can be used as an alternative to proprietary commercial iPSC culture media.

**Fig 1:**
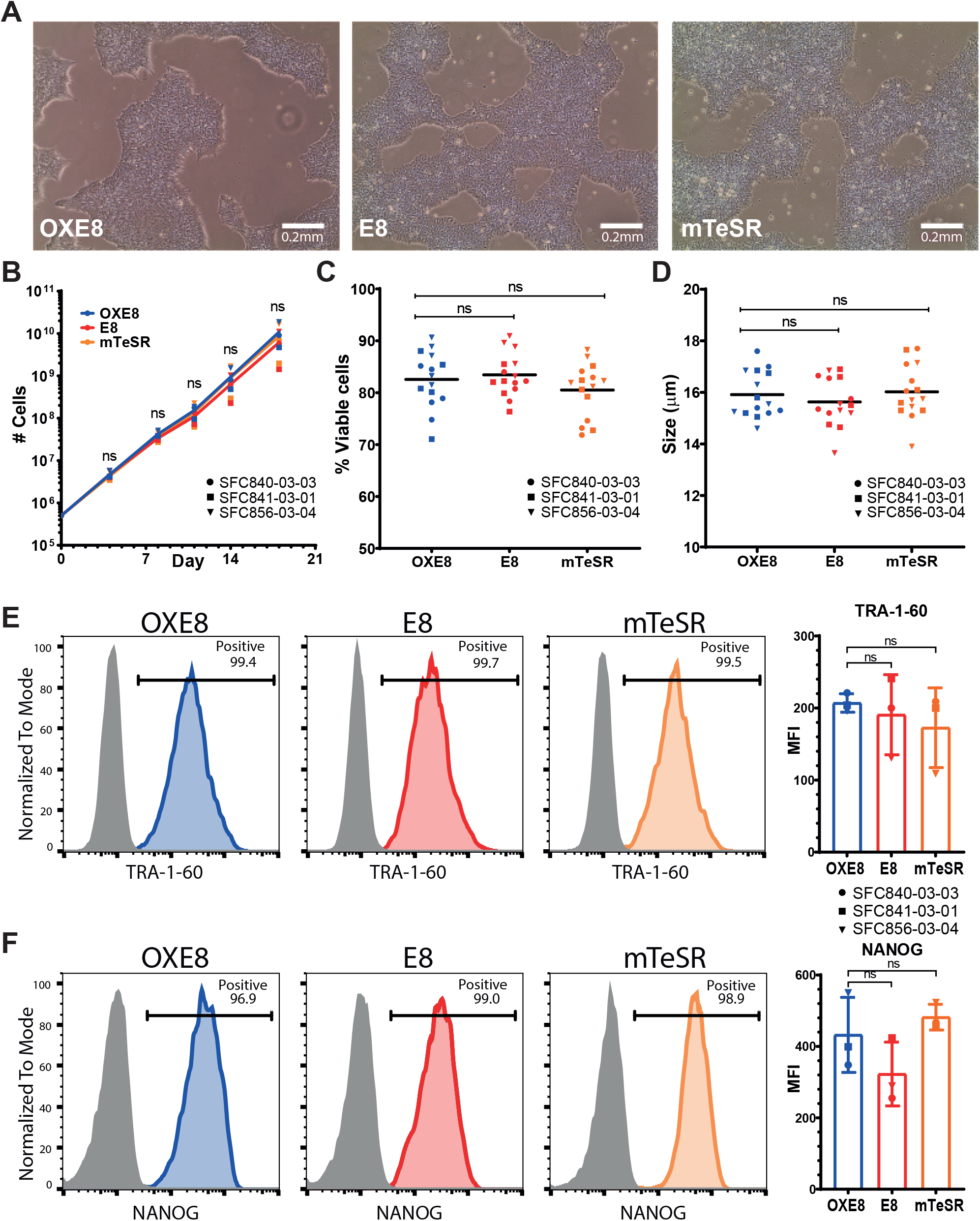
iPSC culture and phenotype. A) iPSC 72h post-seeding at 5×10^4^ cells/cm^2^ in different media. B) Cumulative total number, C) average size, and D) average viability of iPSC over the course of 5 passages. n=3 independent cell lines. Error bars represent ± standard deviation (SD). E) Expression of NANOG or F) TRA-1-60 at passage 5. Histograms show fluorescence intensity (x-axis) normalised to the mode (y-axis) for iPSCs cultured in mTeSR (orange), E8 (red), or OXE8 (blue), relative to the isotype control (grey). Bar chart shows mean ± SD ratio of the geometric mean fluorescence intensity (MFI) compared to the isotype control. n=3 independent cell lines

### XVIVO medium, but not OXM medium, contains undisclosed molecules

We next developed a serum-free, fully defined, open-source medium, named OXM, for differentiation of iPSC to macrophages. Previously we described methodologies for differentiation to macrophages (Karlsson et al., 2008; van Wilgenburg et al., 2013) in which Advanced DMEM or RPMI supplemented with 10% fetal calf serum (FCS), or a serum free alternative, X-VIVO-15™ (XVIVO) were used to culture the cells. However, because the composition of XVIVO is proprietary, we could not know whether the medium contained additives that may affect the phenotype of cells differentiated in this medium. We therefore decided to use aDMEM/F-12 buffered with HEPES as the base of OXM, as used for iPSC culture in OXE8, above. To replace FBS, additional insulin and Tropolone, a cell-permeable iron chelator previously shown to be a suitable substitute for transferrin in cell culture (Field, R.P., Lonza Group, 2003. Animal cell culture. U.S. Patent 6,593,140), were included. M-CSF and IL-3 were supplemented independently.

To identify the cause of any potential differences between XVIVO and OXM-cultured macrophages, we investigated the chemical composition of each medium using high-resolution negative ion liquid chromatography-mass spectrometry (LC-MS). A comparison of the total ion chromatograms (TIC) shows that there are differences in the chemical composition between OXM and XVIVO (Figure 2A) as indicated by the intensity of individual peaks. Each peak was analysed and putatively assigned as described in the supplementary methods. A list of predicted metabolites is given in the supplementary information (Supplementary Table 2). Specifically, we noted a two-fold difference in C6 sugar concentration between XVIVO and OXM. We subsequently independently confirmed this by enzymatic assay, finding 34.3 mM glucose in XVIVO and 16.7 mM in aDMEM/F-12 (Figure 2B). LC-MS also predicted that the level of amino acids were at higher concentration in OXM than in XVIVO except for Tryptophan (predicted concentration: 44 nM in OXM, 6.3 μM in XVIVO)(Supplementary table 2). Additionally, we noted that three peaks in the TIC of the XVIVO media were absent in that of the OXM media (Figure 2A). The best predicted molecules for these peaks are Isolariciresinol 9’-O-alpha-L-arabinofuranoside, (CID: 131751348, Figure 2C), Methyl glucosinolate (CID: 9573942, Figure 2D) and Dithinous acid (CID: 24490, Figure 2E) These molecules were either absent from the OXM media or below detection limit.

**Fig 2:**
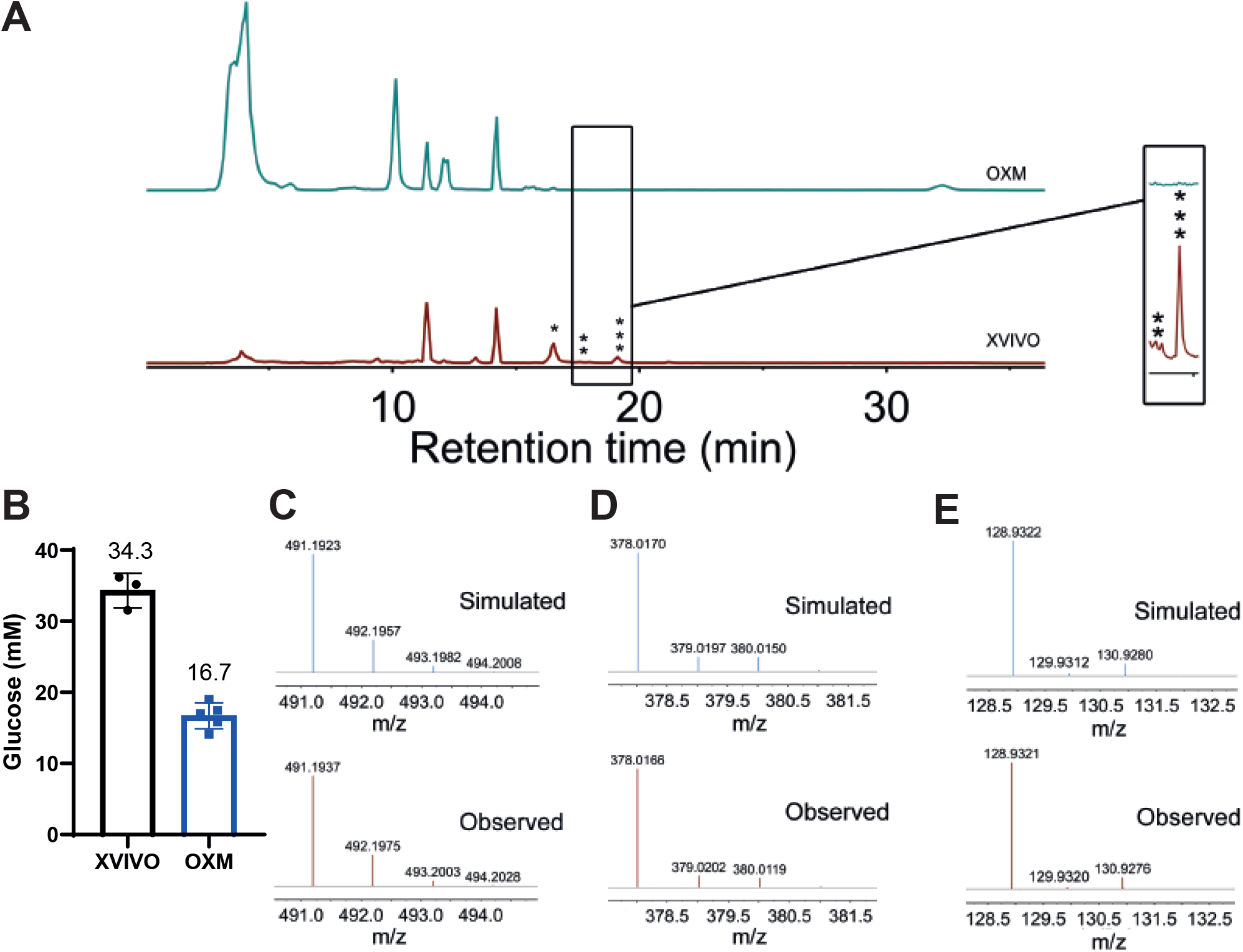
X XVIVO medium, but not OXM medium, contains undisclosed molecules. Negative ion high-resolution LC-MS spectroscopy was used to analyse different compounds in the media. A) Total ion chromatogram (TIC) of the molecules in the XVIVO medium is compared to that of molecules in the OXM medium. B) Concentration of glucose in XVIVO and OXM media. n=3 (XVIVO) and 5 (OXM). Error bars represent ± SD. C) Comparison of the observed MS spectrum of the main peaks absent in OXM with the simulated spectrum of [M-H]^−^ adduct of Isolariciresinol 9’-O-alpha-L-arabinofuranoside (*), D) that of [M+FA-H]^−^ adduct of Methyl glucosinolate (**), and E) that of [M-H]^−^ adduct of Dithionous acid (***).

### Differentiation of iPSC to macrophages in novel fully defined, open source medium, OXM

We next sought to compare the maturation states of OXM-differentiated and cultured macrophages against those differentiated and cultured in XVIVO. As in our earlier methods, differentiation medium was supplemented with IL-3 and M-CSF while, in terminal macrophage differentiation media, IL-3 and tropolone were omitted (Figure 3A, Supplementary table 1). Morphological differences between differentiation cultures were immediately apparent. Embryoid bodies (EBs) produced larger cyst-like structures and greater adherent stromal growth in OXM medium (Figure 3A). By week 3 of differentiation, monocyte-like macrophage precursors cells (PreMacs) had started to be released into the supernatant. This continued for >16 weeks (Figure 3B, 3C). Interestingly, OXM differentiation cultures produced a large number of PreMacs in the first 3 weeks of production before their yield reduced to between 7-8×10^4^ cells/mL/week. XVIVO differentiation cultures were slower to yield PreMacs but by week 6 produced approximately 2-fold more cells, an average of 1.4×10^5^ cells/mL/week (Figure 3B). After 16 weeks the average total yield was (4.48 ± 1.26)×10^7^ cells in OXM, and (7.99 ± 2.61)×10^7^ cells in XVIVO (Figure 3C). PreMacs produced in OXM were significantly smaller than in XVIVO (range 15.8 - 19.5 μm, mean 17.9 μm, vs 13.8 - ≥20 μm, mean 19.5 μm respectively (Figure 3D)). Note that owing to limitations of the Nucleocounter^®^ NC-3000, 20 μm is the upper limit of cell size, so XVIVO-produced cells may have been larger.

**Fig 3:**
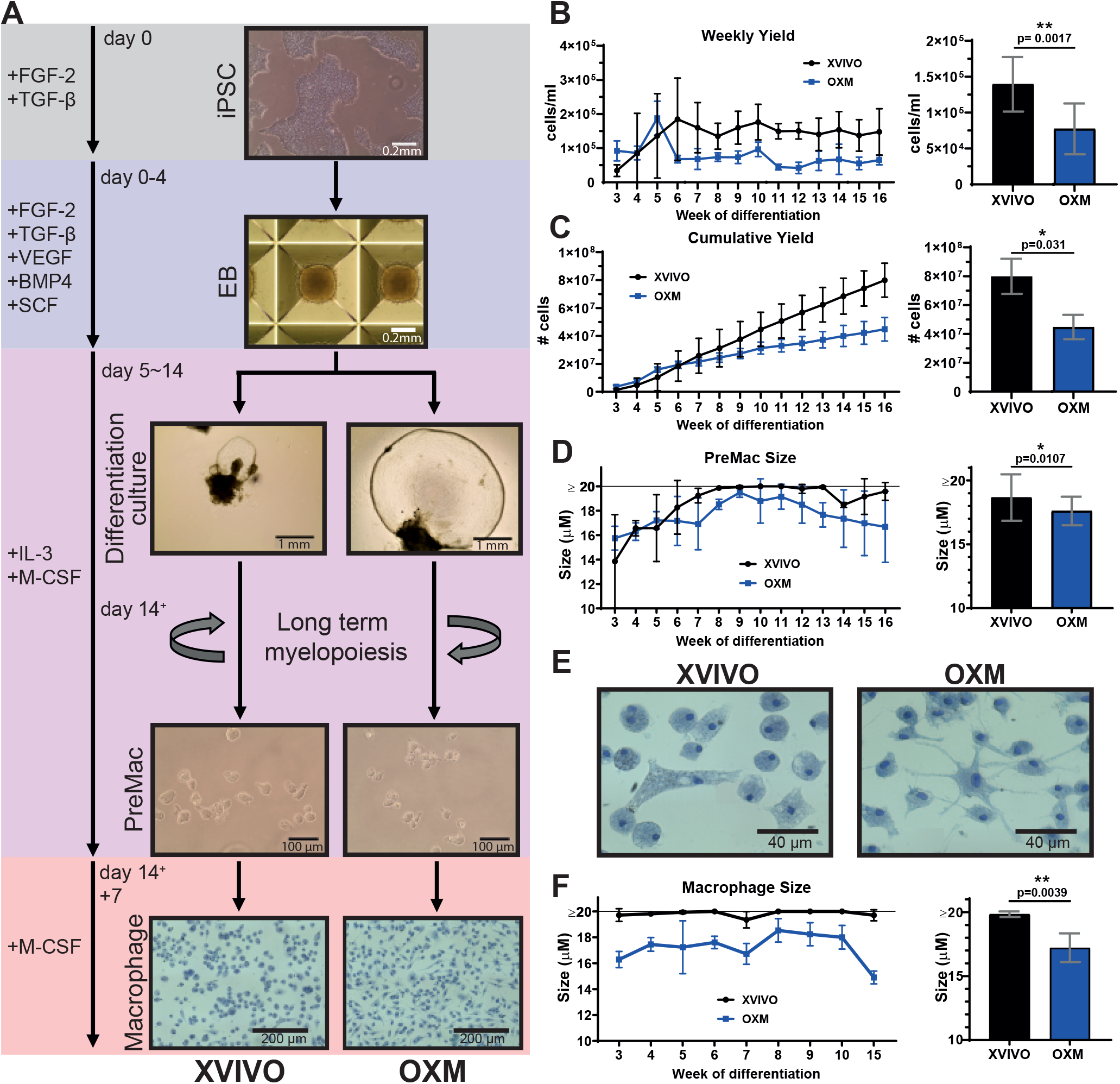
Morphology throughout macrophage differentiation. A) Schematic of iPSC-derived macrophage differentiation. B) Average weekly yield (left) and overall average weekly yield (right) of PreMac cells per mL of media removed from differentiation cultures. C) Cumulative yield of PreMac cells over 16 weeks of differentiation (left), final total yield (right). D) PreMac size over the lifespan of the differentiation culture (left), overall average PreMac size (right). E) Methylene blue (1% w/v) staining to show macrophage morphology. F) Macrophage size over multiple weeks of differentiation (left), overall average macrophage size (right). B-F) XVIVO-cultured cells represented in black vs OXM-cultured cells in blue, mean ± SD. B-C) n=6 across 3 independent cell lines. D, F) n=3 independent cell lines. In all bar charts, significance was calculated by Wilcoxon matched-pairs signed rank t-test. Significance is shown when p<0.05 (*), <0.01 (**), <0.001 (***)

For terminal differentiation of PreMacs into macrophages, cells were cultured a further 7 days in macrophage differentiation medium. After 7 days, as previously observed, XVIVO-cultured cells remain rounded or acquire a bipolar morphology, with large vesicle-like structures visible. OXM-cultured cells were more adherent and morphologically heterogeneous with rounded cells, large flattened cells, and cells with multiple projections (Figure 3A, expanded view in 3E). The size of adherent macrophages was measured after resuspension. OXM-cultured macrophages were significantly smaller than those in XVIVO: mean 17.51 μm (range 16.28 - 18.53 μm) vs mean 19.85 μm (range 19.35 - ≥20 μm), (Figure 3F). In summary, OXM supports the production of more adherent, smaller macrophages than XVIVO, albeit with a lower cumulative yield.

### Macrophages cultured in OXM are phenotypically similar to, but distinct from, XVIVO-cultured cells

To determine macrophage phenotype, surface marker abundance was determined by flow cytometry. Key markers of macrophage lineage, the LPS co-receptor CD14 (Figure 4A), and the pan-leukocyte marker CD45 (Figure 4B), were highly expressed in macrophages derived using both media, with no significant differences. Nor was a difference observed with the expression of scavenger receptor CD163 (Figure 4C). Levels of the Fcγ IgG receptor, CD16, were significantly higher in OXM-cultured cells (Figure 4D), suggesting a more non-classical/intermediate phenotype (Yona and Jung, 2010). Conversely, both the β_2_-integrin receptor, CD11b, and the costimulatory receptor, CD86, were more highly expressed in XVIVO cultured cells, suggesting a more M1-like phenotype (Mosser, 2003) (Figure 4E, 4F). Consistent with our previous reports (Haenseler et al., 2017b; Karlsson et al., 2008; van Wilgenburg et al., 2013), MHC class II (HLA-DR) expression is low in these unstimulated cells in either condition (Figure 4G). Expression of the tissue-resident macrophage marker CD68 was observed intracellularly but not on the cell surface, consistent with prior literature (Gordon and Plüddemann, 2017) (Figure 4H, S1H).

**Figure 4:**
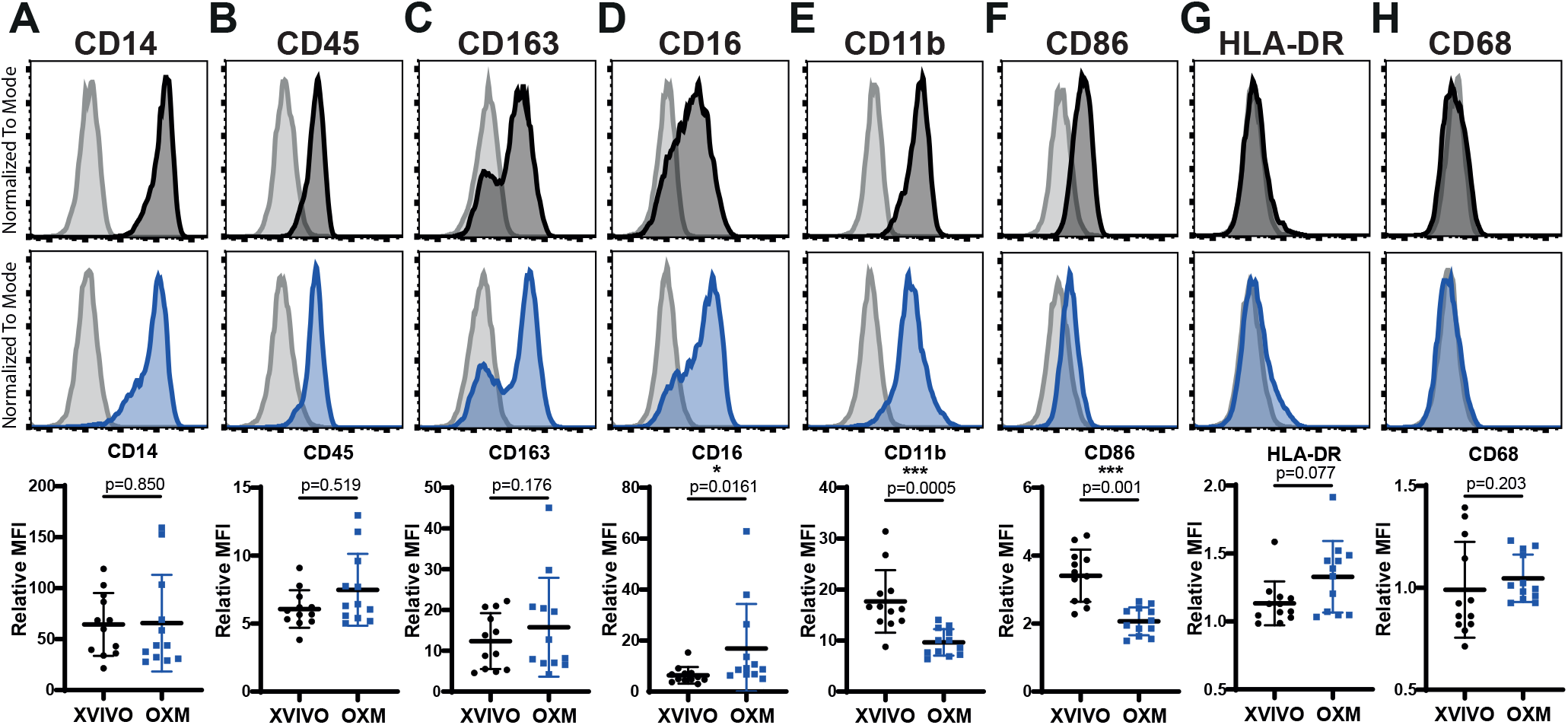
Macrophage surface marker phenotype. Surface expression of A) CD14, B) CD45, C) CD163, D) CD16, E) CD11b, F) CD86, G) HLA-DR, and H) CD68 as measured by flow cytometry on macrophages. Histograms show fluorescence intensity (x-axis) normalised to the mode (y-axis) for macrophages cultured in XVIVO (black) or OXM (Blue), relative to the isotype control (grey). Dot plots show ratio of the geometric MFI compared to the isotype control. Bars display mean ± SD. n=12 across 3 independent cell lines. Significance was calculated by Wilcoxon matched-pairs signed rank t-test. Significance is shown when p<0.05 (*), <0.01 (**), <0.001 (***)

PreMac cells gave broadly similar results to those of fully differentiated macrophages although with differences in CD14 and CD16 expression from full differentiated macrophages (Figure S2A-H). Overall, the PreMacs and macrophages produced using both OXM and XVIVO media display a macrophage phenotype but show small and consistent differences that may reflect differences in polarization.

Finally, to test the phagocytic capacity of macrophages produced under the two conditions, we measured phagocytic uptake of Alexa-488 conjugated zymosan, a yeast-derived particulate glycan. Phagocytic uptake of zymosan after 30 minutes was not significantly different between conditions across 3 genetic backgrounds (Figure S3), indicating that macrophages cultured in OXM are phagocytically competent.

### While transcriptionally very similar, OXM-cultured macrophages have a more homeostatic, and XVIVO-cultured macrophages a more immunologically active, signature

We used RNA-Seq to investigate the detailed expression profile of the iPSC-derived macrophages. We first compared the macrophages cultured in XVIVO or OXM to a published dataset of iPSC-derived human microglia (tissue resident macrophages of the brain) that had been differentiated via Hematopoietic Progenitor Cells (iHPCs) derived using modified protocols from prior reports (Abud et al., 2017; Kennedy and Keller, 2007; Sturgeon et al., 2014). PCA analysis, in which PC1 explained 27% of the variance, and PC2 explained 18%, showed tight clustering of OXM- and XVIVO-cultured cells together. The iPSC-derived macrophages showed closest similarity to microglia, and least similarity to pluripotent cells and polarized, definitive lineages such as dendritic cells (Figure 5A). Further differential expression analysis found 15,951 genes shared in both OXM and XVIVO populations, and 2335 genes unique to either OXM (1331) or XVIVO (1004) (Figure 5B). Analysis of the sources of this variation showed that the media used explained approximately 30% of the variation between samples, while sequential harvests of cells from the differentiation cultures accounted for approximately 15%, and the remainder was due to residual, undefined sources (Figure 5C).

**Figure 5:**
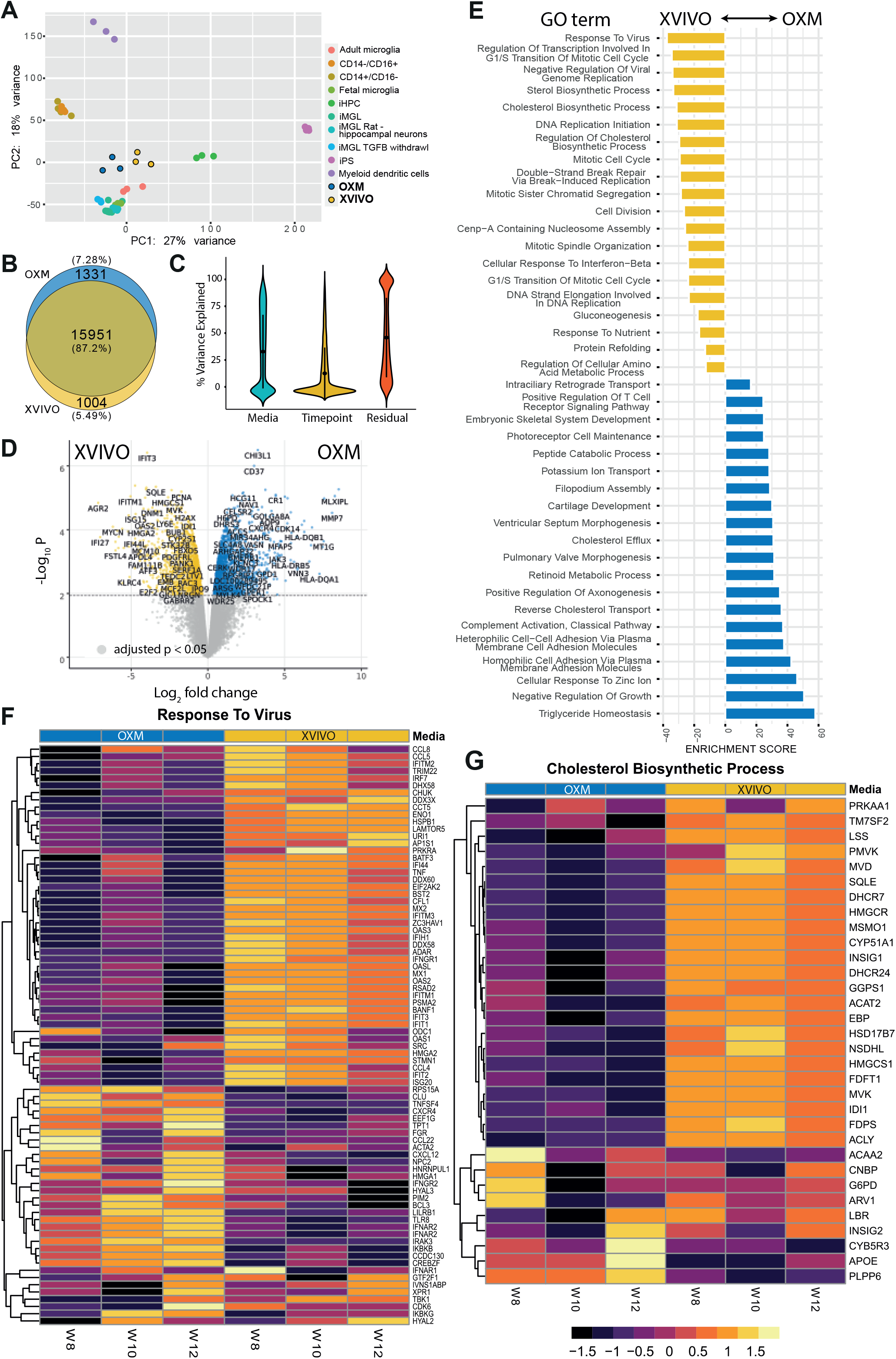
Transcriptome of iPSC-derived macrophages. Transcriptome of macrophages as measured by RNA-Seq of 3 biological replicates per media condition from one genetic background (SFC840-03-03). A) PCA plot of transcriptional profile for iPSC-derived macrophages cultured in XVIVO or OXM compared to iPSC, induced haematopoietic progenitor cells (iHPC), microglia (adult, fetal, iPSC-derived, and iPSC-derived with TFGB withdrawal), monocytes, and dendritic cells previously reported (Abud et al., 2017). B) Venn diagram representing total number of genes expressed by iPSC-derived macrophages in either media composition. C) Violin plot of the percentage variance attributed to each variable (media, timepoint, and residual). D) Volcano plot of differentially expressed genes between the media compositions. y=log_10_ (P value), x=log_2_ (fold change). Yellow represents genes with lower expression in OXM (higher expression in XVIVO) and blue represents higher expression in OXM (lower in XVIVO). Genes were considered differentially expressed if p<0.05 E) GO term enrichment between media compositions. Top 15 most enriched results in each media are shown. Yellow represents terms enriched in XVIVO-cultured cells, and blue in OXM-cultured cells. F-G) Heatmaps of GO terms for; “Response to Virus” and “Cholesterol Biosynthetic Process”. Results across 3 repeat measurements are shown. XVIVO in yellow, OXM in blue. The colour in each row represents the z-score of the log_2_ fold difference from the mean TPM value for that gene.

The three genes most significantly upregulated in OXM versus XVIVO were the carbohydrate binding lectin, *CHI3L1*, the tetraspanin, *CD37*, and the protease, *HTRA4*. In XVIVO the most upregulated genes were the interferon-inducible protein *IFIT3*, the endonuclease *RNASE1*, and the epoxidase *SQLE* (Figure 5D). The most enriched GO term in XVIVO compared to OXM-cultured cells, was “Response to virus” (Figure 5E, 5F). Multiple interferon inducible genes, e.g. *MX2*, *RSAD2*, and the *IFIT* family of proteins, were more highly expressed in XVIVO-cultured cells. However, some antiviral receptors including *TLR8*, *IFNAR2*, and *INFGR2*, are more highly expressed in OXM-cultured cells. We also found enrichment of metabolism-related processes. In OXM-cultured cells, the most highly enriched GO term was “triglyceride homeostasis” (Figure S4A), and in XVIVO-cultured cells the cholesterol biosynthetic process was highly enriched with 23/32 genes in this term upregulated in these cells (Figure 5E, 5G). The term “DNA replication initiation” was enriched in XVIVO-cultured cells (Figure S4B). To test whether XVIVO cells may therefore be in cell cycle, we measured total KI67, which transcriptionally has 2.66-fold higher expression in XVIVO-cultured cells Figure S4C), by immuno-staining and flow cytometry. This confirmed that expression is higher, but note that only a small proportion of cells (XVIVO; 1.15% +/− 0.90%, OXM; 0.54% +/− 0.55%) are in cell cycle (Figure S4D). In OXM, the inflammation related term “complement activation, classical pathway”, cell adhesion, and several developmental terms are enriched (Figure 5E, S4E). The enrichment of the cell adhesion term supports visual observations that OXM cultured cells were more adherent to the culture plate than XVIVO cultured cells. Overall, transcriptomic analysis has shown that although cells cultured in these different media are highly similar, they display a greater tendency towards proliferation and an activated antiviral response in XVIVO, and towards homeostasis, adhesion, and inflammation in OXM.

### OXM and XVIVO cultured macrophages have different classical and alternative activation profiles

To test whether the subtle transcriptomic differences between the cells differentiated in the two media resulted in functional detectable differences, we measured cytokine secretion and polarisation after stimulation. Cytokine secretion was measured at resting state and after classical (LPS and INFγ) or alternative (IL-4) activation, as previously described (Jiang et al., 2012; van Wilgenburg et al., 2013). Cytokine profiles under both conditions were consistent with the previous reports (Figure 6A). OXM-cultured cells produced noticeably more GROα, ICAM-1, IL-1ra, and IL-8 in the resting state, while XVIVO-cultured cells produced more IP-10 (CXCL10). After classical activation, XVIVO-cultured cells produced more IL-10, IL-12p70, and TNFα, while OXM-cultured cells produced more IL-1β and RANTES (CCL5). Alternative activation suppressed secretion of resting state cytokines, and increased secretion of MIF in both media, but increased secretion of Serpin E1 only in OXM-cultured cells. Consistent with the results above, LPS-induced TNFα secretion from OXM-cultured cells quantified by enzyme-linked immunosorbent assay (ELISA) was lower than from XVIVO cells (Figure 6B).

**Figure 6:**
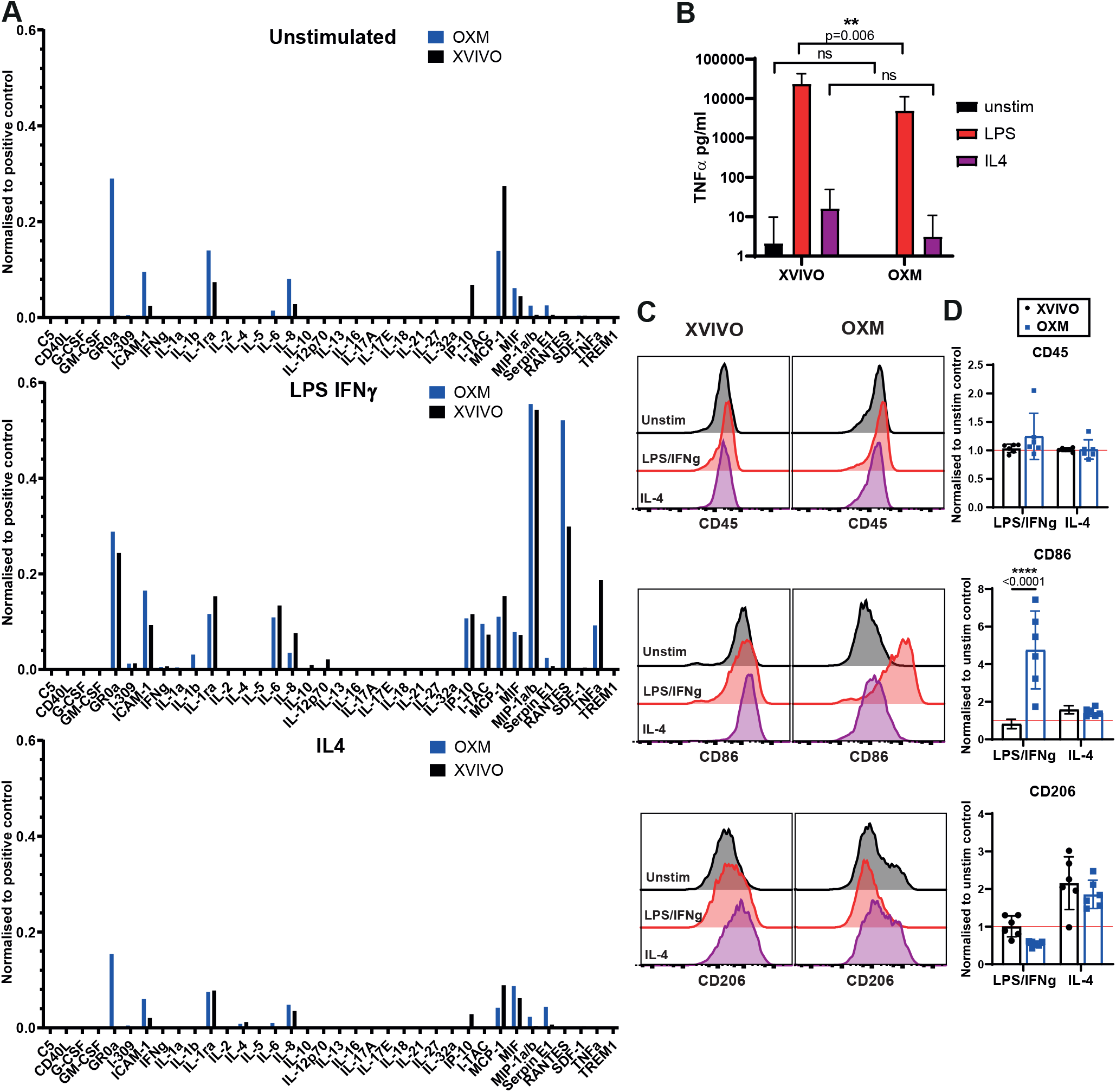
Macrophage activation states between culture media. A) Cytokine secretion into the supernatant under unstimulated resting conditions (top), stimulated with 100 ng/mL LPS + 20 ng/mL IFNγ (middle), or 50 ng/mL IL-4 (bottom) normalised to a positive control. n=1. B) TNFα secretion from unstimulated (black), 100 ng/mL LPS (red), or 50 ng/mL IL-4 (purple) stimulated cells. Mean ± SD, n=14 in 3 independent cell lines. D) Surface expression of CD45, CD86, or CD206 measured by flow cytometry after stimulation, coloured according to (B). Histograms show fluorescence intensity (x-axis) normalised to the mode (y-axis). E) Geometric MFI relative to unstimulated control. Mean ± SD, n=6 in 2 independent cell lines. B-E) Significance was calculated by 2-way ANOVA, Sidak’s multiple comparison test. Significance is shown when p<0.05 (*), <0.01 (**), <0.001 (***), <0.0001 (****)

Considering the differences in cytokine production, we next assessed macrophage polarisation towards classically defined M1 and M2 states after activation. This was measured by immuno-staining for CD86 and CD206 respectively (Mosser, 2003; Roszer, 2015) (Figure 6C-D). Although activation had no effect on CD45 expression in both conditions, CD86 expression significantly increased in OXM-cultured cells compared to XVIVO-cultured cells after classical activation (4.75-fold vs. 0.81-fold). CD206 expression increased in both conditions after alternative activation and was not significantly different between conditions (1.86-fold vs 2.16-fold) (Figure 6C-D). In conclusion, although cells cultured under either condition can respond well to stimuli, they differ in their cytokine profile, and cells cultured in OXM have a greater capacity for polarization than those in XVIVO.

### OXM-cultured macrophages are more susceptible to HIV-1 infection, but not to Zika virus

Due to their positioning as sentinel cells throughout the body, tissue resident macrophages are often the first members of the immune system to respond to infection. Consequently, several pathogens have evolved to use these cells as a host during early infection, including viruses like human immunodeficiency virus-1 (HIV-1) and Zika virus (ZIKV). In the case of HIV-1, macrophages are hypothesised to act as a potential reservoir for latent infection due to the long lived, non-replicating state of these cells (Honeycutt et al., 2016, 2017). To determine whether iPSC-macrophages would be a suitable model for modelling this disease, macrophages were infected with a macrophage-tropic (CCR5-tropic Ba-L envelope), GFP expressing, HIV-1 pseudotype virus. Infectivity was measured by expression of GFP after 72 hours by flow cytometry. A significantly higher proportion of macrophages cultured in OXM than in XVIVO were infected (6.76 ± 3.69% compared to 1.77 ± 1.23%; See Figure 7A). Expression of the HIV-1 entry receptors CD4 and CXCR4 was not-significantly different between conditions, and CCR5 was higher in XVIVO-cultured macrophages (MFI relative to isotype: 6.16 ± 3.18 XVIVO vs 2.67 ± 1.52 OXM; Figure 7B). This difference in viral entry may be explained by the higher expression of known HIV-1 restriction factors in XVIVO-cultured cells including the *IFITM* and the *APOBEC* families of proteins, and *TRIM5*α (Figure 5F). To test whether differences in susceptibility are restricted to lentiviruses, we also assessed replication ZIKV strain MR-766 (Uganda, 1947) in these cells. Cells produced in both media released virus after an eclipse phase of about 6 hours post infection with no difference in output titre ((3.36 ± 1.83)×10^7^ pfu/mL XVIVO, (2.26 ± 1.14)×10^7^ pfu/mL OXM at 48hpi; see Figure 7C). Therefore OXM-cultured macrophages are a good model for HIV-1 infection, and differences in viral susceptibility between culturing conditions are not pan-viral.

**Figure 7:**
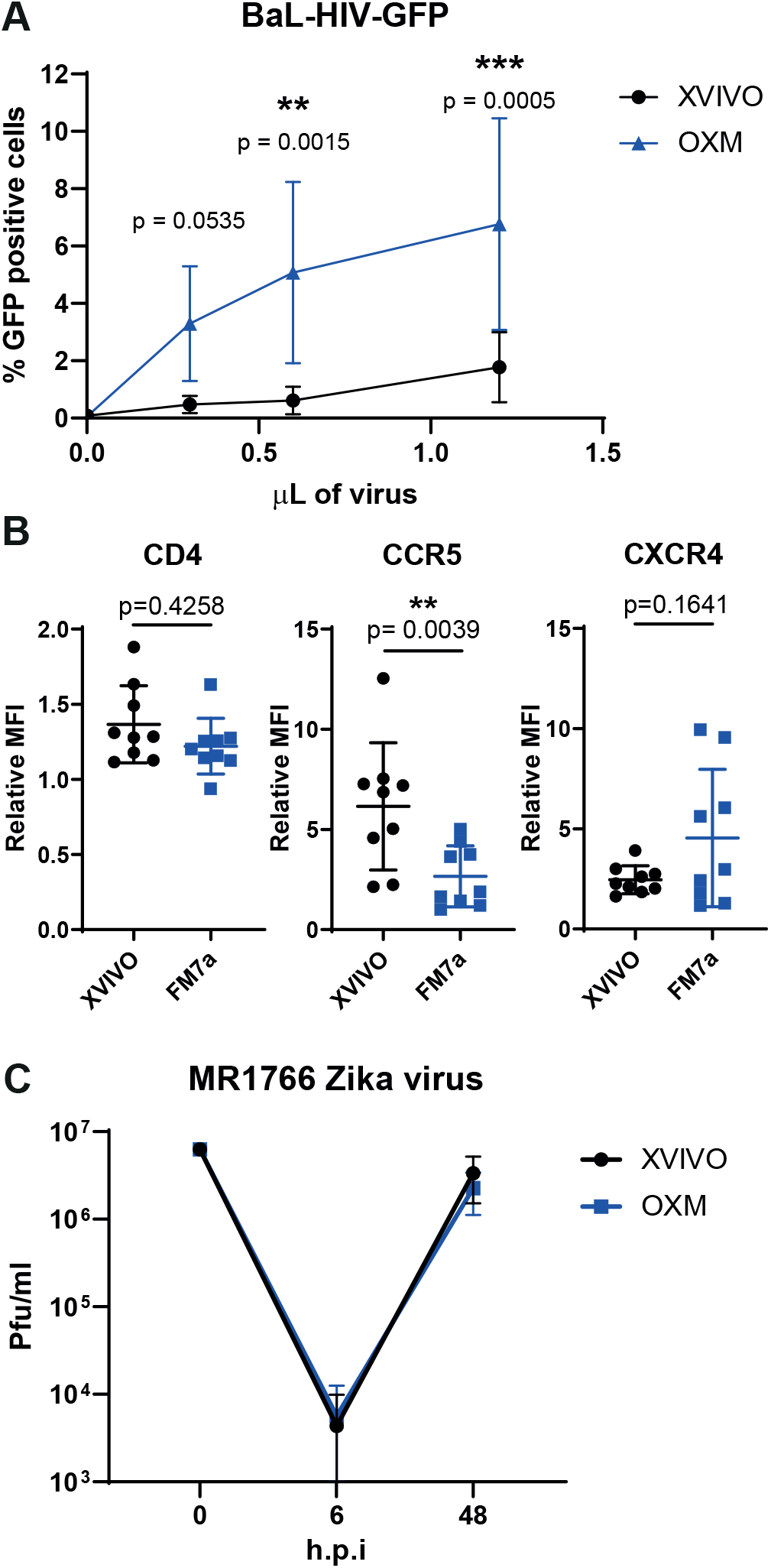
Macrophage infection by HIV-1 or Zika virus in different culture conditions. A) Percentage of macrophages infected with a GFP-encoding, single-round HIV-1 pseudotyped virus measured by flow cytometry. B) CD4, CCR5, and CXCR4 surface expression measured by flow cytometry shown as ratio of geometric MFI to the isotype control. C) MR1766 Zika virus titre from infected macrophages shown in plaque forming units (pfu)/ml. Significance was calculated by A) 2-way ANOVA, Sidak’s multiple comparison test, B) Wilcoxon matched-pairs signed rank t-test. A-C) Mean ± SD n=8 in 2 independent cell lines. Significance is shown when p<0.05 (*), <0.01 (**), <0.001 (***)

## Discussion

Here we have established two fully defined media for the growth of iPSC and iPSC-derived macrophages based on aDMEM/F-12 called OXE8 and OXM, respectively. Both media are easy to make and comparatively cheaper than commercial media, with OXE8 approximately 1/5 the price of commercial E8, and OXM 1/3 the cost of XVIVO before supplementation with M-CSF and IL-3. We observed no significant difference between hiPSC grown in OXE8 medium and in current, commercial alternatives. We did observe subtle but significant differences in yield, morphology, and phenotype of iPSC-derived macrophages cultured in OXM compared to the commercial alternative, XVIVO. OXM offers an alternative to XVIVO, and avoids the presence of several, undisclosed, supplements in the latter medium.

In order to culture iPSC, researchers are generally reliant on commercial media that are not only proprietary, but also very expensive, limiting the ability of many laboratories to undertake iPSC-based research. There has been a recent push to lower this cost (Kuo et al., 2020). The alternative that we describe here is also cheaper than the commercial alternatives. We have shown that culturing in OXE8 does not adversely affect iPSC growth or pluripotent state over numerous passages.

The need to define a fully open-source medium for differentiating macrophages from iPSC was driven by two requirements. Firstly, to understand the finer changes in macrophage response to stimuli we must have a clear understanding of the resting conditions of these cells. Secondly, LC-MS analysis indicated the potential presence of several undisclosed molecules in the commercial serum-free alternative media, XVIVO™ 15, some of which could inhibit, or polarize the macrophage response if present. Our analysis putatively identified Isolariciresinol 9’-O-alpha-L-arabinofuranoside, a lignan glycoside, which is known to have anti-inflammatory effects (Liu et al., 2018) as well as methyl glucosinolate and dithionous acid. While the precise functions of these molecules are unknown, it is worth noting that indole glucosinolates have been reported to have potent anti-inflammatory effects (Vo et al., 2014). In its ionised form (dithionite), dithionous acid is a strong reducing agent capable of affecting the redox state of haem-containing enzymes, such as cytochrome b, which may affect cell physiology (Berton et al., 1986).

We found that basing our new media, OXM, on aDMEM/F-12, did not prevent differentiation of macrophages from iPSC. However, we did see considerable differences in both yield and morphology. OXM-cultured cells appeared to be more adherent, with the increased expression of adherence-related genes. It is possible that the decreased yield could be due to increased adherence of PreMac cells to the tissue culture plastic in differentiation cultures, which limited the harvestability of the cells to once per week. We have previously shown that XVIVO-cultured iPSC-macrophages are significantly larger than blood monocyte-derived macrophages (van Wilgenburg et al., 2013), which are closer in size to OXM-cultured cells. Considering the very high levels of glucose in XVIVO medium, we suspected that there could be major differences in cell metabolism. We were therefore unsurprised to see that metabolism-related GO terms were among the most abundant terms differentially enriched between the media types. Whereas OXM-cultured cells were enriched for triglyceride homeostasis, reverse cholesterol transport and peptide catabolic process, XVIVO-cultured cells upregulated many genes involved in lipid metabolism and cholesterol biosynthesis such as fatty acid synthase (*FASN*), 7-Dehydrocholesterol Reductase (*DHCR7*), and Acetyl-CoA Acetyltransferase 2 (*ACAT2*).

Macrophage metabolism differs substantially between activated states (M1 vs M2), with fatty acid synthesis upregulated in M1 cells and fatty acid oxidation upregulated in M2 cells (Caputa et al., 2019). Unsurprisingly then, we observed that XVIVO-cultured cells had higher expression of markers of classical M1 (pro-inflammatory) polarization (Boyette et al., 2017; Mosser, 2003). It is possible that this activated state in XVIVO is responsible for their relative insensitivity to LPS and IFNγ – induced changes in CD86 expression, unlike OXM-cultured cells which displayed a stronger potential to polarize, although it does not appear to impact the cytokine response by these cells. Conversely, OXM-cultured cells showed similarities to non-classical inflammatory monocytes, with increased expression of CD16 and the constitutive secretion of some inflammatory cytokines such as IL-6 and GROα. This cytokine profile observed is consistent with our previous work (Haenseler et al., 2017b) in which iPSC-derived microglia released similar cytokines in monoculture that were reduced when co-cultured with neurons. This suggests that tissue resident macrophages likely receive anti-inflammatory signals from neighboring stromal cells as part of a regulatory feedback system. Overall, while they did not clearly fall into a conventional phenotype, such as “classical”, “non-classical”, “M1” or “M2”, such categories, which are largely derived from work with blood monocyte-derived macrophages cultured in FBS-containing medium (Murray et al., 2014; Nahrendorf and Swirski, 2016), are an imperfect system for the phenotypic characterization of tissue resident macrophages.

One surprising observation was that XVIVO-cultured cells appear to be transcriptionally primed for an antiviral response. We found higher expression of many interferon inducible genes and constitutive secretion of the interferon inducible cytokine IP-10 (CXCL10). This clearly translated into a reduced infectivity of HIV-1 pseudotype virus in these cells compared to OXM-cultured cells despite the cells having higher expression of the surface receptor CCR5. However, it did not result in a difference in ZIKV infection of these cells. This is supported by the fact that OXM-cultured cells had higher expression of the interferon receptors *IFNAR2* and *IFNGR2*, and therefore are likely capable of mounting a strong antiviral response. It is worth noting that although expression of interferon genes was lower in OXM-cultured cells, it was not absent. These cells may therefore be a better model of pre-viral exposure macrophages.

To conclude, we describe the successful development of low-cost, defined, and open-source media for culturing human iPSC and differentiating them into macrophages: OXE8 and OXM, respectively. We hope that the open-source nature of these media will enable them to be used as foundations for future improved differentiation protocols not only for macrophages, but also other iPSC-derived cell types.

## Experimental Procedures

### iPSC lines

The derivation and characterisation of the iPSC lines used in this study is described elsewhere: SFC840-03-03 (Fernandes et al., 2016), SFC841-03-01 (Dafinca et al., 2016) SFC856-03-04 (Haenseler et al., 2017b)). See Supplemental Information for more information.

### iPSC culture

iPSC were cultured in their stated media (Supplementary table 1) on Geltrex™ (Gibco, #A1413201)-coated tissue culture dishes and passaged using TrypLE™ Express (Gibco, #12604013). For 24 hours after plating, media was supplemented with 10 μM Y-27632 (Abcam, ab120129). Cells were incubated at 37°C, 5% CO_2_. See Supplemental Information for more information.

### Macrophage differentiation

iPSC were differentiated into macrophages using a protocol based on that previously described (van Wilgenburg et al., 2013). The updated method is as follows. Aggrewell™ 800 plates (Stemcell technologies, #34815) were prepared by addition of 0.5 mL Anti-Adherence Rinsing solution (Stemcell technologies, #07010) and centrifugation at 3000g for 3 minutes to remove bubbles from the microwells. Rinse solution was then aspirated and replaced with 1 mL of 2X concentrated EB medium (1X EB media: OXE8 medium supplemented with 50 ng/mL BMP4 (Peprotech, #PHC9534), 50 ng/mL VEGF (Peprotech, #PHC9394), and 20 ng/mL SCF (Miltenyi Biotec, #130-096-695) supplemented with 10 μM Y-27632. iPSC were resuspended by washing with PBS, incubating in TrypLE Express for 3-5 minutes at 37°C, 5% CO_2_, followed by gentle lifting in Advanced-DMEM/F12 to achieve single cell suspension. Cells were counted and pelleted by centrifugation at 400g for 5 minutes. After centrifugation, cells were resuspended at 4×10^6^ cells/mL in OXE8 supplemented with 10 μM Y-27632 and 1 mL added to the Aggrewell. The Aggrewell plate was then spun at 100g for 3 minutes with no braking to encourage even distribution of cells across microwells. Cells were incubated for 4 days at 37°C, 5% CO_2_ with daily feeding of 75% media change with EB medium, by aspiration of 1 mL by pipette and gentle addition of 1 mL fresh media twice to avoid disturbance to the microwells. After 4 days, EBs were lifted from the plate using a Pasteur pipette and passed over a 40 μm cell strainer to remove dead cells, before washing into tissue culture plate with differentiation media (Supplementary table 1). EBs were divided evenly into 2 T175 flasks and topped up to 20 mL with differentiation medium. Differentiation cultures were incubated at 37°C, 5% CO_2_, with weekly feeding of an additional 10 mL until macrophage precursors (PreMac) cells started to be produced. After this point PreMac cells were collected weekly and a minimum of equal volumes of media to the volume removed were replaced into the differentiation cultures. Each harvest involved a 25-50% media change. Harvested cells could be stored or transported on ice for up to 48h without loss of viability, and were either used directly or plated in appropriately sized tissue culture plates for further culturing in XVIVO or OXM Macrophage medium (Supplementary table 1) for a further 7 days, with a 50% media change on day 4.

### Cell count, size, and viability measurements

Cell count, size, and viability were measured using the Nucleocounter^®^ NC-3000 (Chemometec) after staining for live/dead cells using Solution-13 AO-DAPI stain (Chemometec, #910-3013). See Supplemental Information for more details.

### Flow Cytometry

Cells were stained directly without fixation in FACS buffer (PBS supplemented to 1% FBS, 10 μg/mL human-IgG (Sigma, #I8640-100MG), and 0.01% Sodium azide) and compared to isotype controls with the same fluorophores from the same company. Fluorescence was measured using the BD LSRFortessa™ X-20 (BD Biosciences) and analysed on FlowJo version 10. See Supplemental Information for more details.

### Cytokine and Chemokine release

Cytokine and Chemokine secretion was assessed as previously described (van Wilgenburg et al., 2013) using the Proteome Profiler Human Cytokine Array Kit (R&D systems, #ARY005B). TNFα was quantified by ELISA (Invitrogen, 88-7346-88) according to manufacturer’s instructions. See Supplemental Information for more details.

### Phagocytosis

Phagocytosis was measured by uptake of Zymosan A (*S. cerevisiae*) BioParticles™ Alexa Fluor-488 (ThermoFisher, #Z23373) after 30 minutes. Uptake was measured by measurement of fluorescence using the BD LSRFortessa™ X-20. See Supplemental Information for more details.

### HIV-1 infection Assay

7-day differentiated macrophages were infected for 72 hours with a GFP expressing HIV-1 lentiviral vector pseudotyped with strain Ba-L envelope virus diluted in macrophage medium. Percentage infected cells was determined by measurement of fluorescence using the BD LSRFortessa™ X-20 flow cytometer. See Supplemental Information for more details.

### Zika virus infection assay

Macrophages were infected with 2 × 10^6^ pfu of ZIKV isolate MR1766 for 4 hours before replacing with fresh medium. Supernatant was collected at stated time points and stored for later quantification of virus by plaque assaying on Vero-76 cells. See Supplemental Information for more details.

### RNA-Seq

RNA was extracted from 1×10^6^ macrophages cultured in a 6-well plate, differentiated from PreMac cells harvested at week 8, 10, and 12 of differentiation, and sequenced by Novogene. Results were analysed in-house. See Supplemental Information for more details.

### Liquid chromatography-mass spectrometry (LC-MS) analysis of media

Samples of the media were prepared for LC-MS analysis as described before (Ebrahimi et al., 2020). See Supplemental Information for more details.

### Quantification of glucose concentration

Glucose concentration in the media was quantified using a modified protocol for the Glucose (HK) Assay Kit (Supelco, # GAHK20-1KT). See Supplemental Information for more details.

### Statistical analysis

All statistical analysis used is reported in the figure legends and was carried out using GraphPad Prism version 8.2.1 or R version 4.0. In all cases data was considered significant when p<0.05.

## Acknowledgements

This project was principally funded by a Wellcome Trust Four-year PhD Studentship. Liquid chromatography-mass spectrometry samples were kindly run in the facility of Prof. James McCullagh (Department of Chemistry). The work in the group of Prof. James McCullagh is supported by Wellcome Institutional Strategic Support Fund. Stem cell work was carried out in the James Martin Stem Cell Facility which has received financial support from the Oxford Martin School.

## Author Contributions

Conceptualization, A.V-J, S.A.C, and W.S.J. Methodology, A.V-J and W.S.J. Investigation, A.V-J, K.H.E, C.B, E.P, J.G-J, and W.S.J. Data Curation, S.S. Formal Analysis, A.V-J, S.S and K.H.E. Visualization, A.V-J, S.S, K.H.E, and P.K.R. Supervision, P.K.R and E.P. Writing – Original Draft, A.V-J, K.H.E, and W.S.J. Writing – Review & Editing, A.V-J, S.A.C, and W.S.J. Funding Acquisition, A.V-J, S.A.C and W.S.J.

