## Supplementary Tables and Information for "Differentiation of Human induced Pluripotent Stem Cells to Authentic Macrophages using Fully Defined, Serum Free, Open Source Media"

**Supplementary historical perspective related to the Introduction**

We have self-consciously built on the methodology for tissue culture established by Harry Eagle in the 1950s (Eagle, 1955b, 1955a, 1959). Dulbecco’s modification of MEM (DMEM) involved increasing the concentration of all amino acids and vitamins by 4-fold and produced significant increases in cell and virus yield (Dulbecco and Freeman, 1959; Smith et al., 1960). Because of the variabily of quality of serum products, and their undefined composition, Puck and Ham developed the “F” series of increasingly complex basal media, in which serum concentration could be progressively reduced by the addition of fatty acids and organic amines, culminating in one, called F12, that would support the growth of certain cell lines without serum (Ham, 1965; Ham and McKeehan, 1979). De Vallis and Sato went on to uncover most of fetal bovine serum’s (FBS) key components so that it could be replaced with fully synthetic ingredients (Bottenstein and Sato, 1979; Hayashi and Sato, 1976; Murakami et al., 1982; Orly and Sato, 1979) (Barnes and Sato, 1980). From this period onwards, serum-free, and serum-reduced media were typically based on the 1:1 mixture of DMEM and Ham’s F12, (Bottenstein and Sato, 1979) supplemented by insulin and transferrin, together with trace elements such as selenium (McKeehan et al., 1976). For convenience, the most common supplements including transferrin, insulin, selenium, vanadium, ethanolamine, L-ascorbic acid, and glutathione are present in defined, open-source commercial formulations such as “advanced” DMEM/F12.

A synthetic approach was successfully applied to the culture of human pluripotent stem cells by the Thompson group, culminating in E8 medium, in which the only hormonal supplements required are insulin, FGF-2, and TGF-β (Chen et al., 2011), and attachment is provided by vitronectin. A further, chemically-defined derivative of E8 medium has recently been reported (Kuo et al., 2020), and commercial, proprietary variants of E8 are available (Silpa et al.)

**Supplemental Figure legends**

**Fig S1: Karyotype of long-term cultured iPSC in OXE8. Related to Figure 1.**

A) Diagram of copy number variant (CNV) as compared to the human genome GRCh38 in 5 different iPSC lines cultured in OXE8 media ≥3 passages. Double allele deletions are shown in red, single allele deletions in orange, single allele duplications in blue, and double allele duplications in purple. Where copy number was between 1.5 and 2.5 sequence was not annotated, except in the sex chromosomes where it is displayed in green. B) . Analysis was carried out on GenomeStudio (Illumina) version 2.0 allowing for detection of extended homozygosity.

**Figure S2: Macrophage total marker phenotype. Related to Figure 4.**

Total expression of A) CD14, B) CD45, C) CD163, D) CD-16, E) CD11b, F) CD86, G) HLA-DR, and H) CD68 as measured by flow cytometry in macrophages. Histograms show fluorescence intensity (x-axis) normalised to the mode (y-axis) for macrophages cultured in XVIVO (black) or OXM (Blue), relative to the isotype control (grey). Dot plots show ratio of the geometric MFI compared to the isotype control. Bars display mean ± SD. n=12 across 3 independent cell lines. Significance was calculated by Wilcoxon matched-pairs signed rank t-test. Significance is shown when p<0.05 (*), <0.01 (**), <0.001 (***)

**Figure S3: PreMac surface marker phenotype. Related to Figure 4.**

Surface expression of A) CD14, B) CD45, C) CD163, D) CD-16, E) CD11b, F) CD86, G) HLA-DR, and H) CD68 as measured by flow cytometry on PreMac cells. Histograms show fluorescence intensity (x-axis) normalised to the mode (y-axis) for macrophages cultured in XVIVO (black) or OXM (Blue), relative to the isotype control (grey). Dot plots show ratio of the geometric MFI compared to the isotype control. Bars display mean ± SD. n=12 across 3 independent cell lines. Significance was calculated by Wilcoxon matched-pairs signed rank t-test. Significance is shown when p<0.05 (*), <0.01 (**), <0.001 (***)

**Figure S4: Phagocytosis in different culture conditions. Related to Figure 4.**

Number of macrophages that are positive for Alexa-488 conjugated zymosan coated beads after 30-minute incubation, as measured by flow cytometry. Control cells were pre-treated with 10µM Cytochalasin D for 1 hour to inhibit phagocytosis. n=8 across 3 independent cell lines. Significance was calculated by 2-way ANOVA, Sidak’s multiple comparison test. Significance is shown when p<0.05 (*), <0.01 (**), <0.001 (***)

**Figure S5: Macrophage cell cycle state. Related to Figure 5.**

A-B, E) Heatmaps of GO terms for; A) Triglyceride Homeostasis, B) DNA Replication Initiation, and E) Homophilic Cell Adhesion Via Plasma Membrane Adhesion Molecules. Results across 3 repeat measurements are shown. XVIVO in yellow, OXM in blue. The colour in each row represents the z-score of the log_2_ fold difference from the mean TPM value for that gene. C) Mean TPM values for *MKI67* (KI67) expression in XVIVO vs OXM-cultured cells. D) Percentage of KI67 positive cells measured by immune-staining and flow cytometry. XVIVO in black, OXM in blue. n=6 in 2 independent cell lines. Mean and standard deviation shown. Significance was calculated by Wilcoxon matched-pairs signed rank t-test. Significance is shown when p<0.05 (*), <0.01 (**), <0.001 (***), <0.0001 (****)

**Supplemental Table Legends**

**Supplementary Table 1: iPSC and macrophage culture media. Related to Figure 1 and 3.**

| **Media** | **Component** | **Final Concentration** | **Supplier** | **Catalogue number** |
| --- | --- | --- | --- | --- |
| **OXE8** | Advanced DMEM/F-12 | 97.4% | Gibco | 12634010 |
|  | GlutaMAX (100x) | 2mM | Gibco | 35050-038 |
|  | Heparin solution 0.2% | 100ng/mL | STEMCELL^TM^ | 07980 |
|  | Ascorbic Acid-2-phosphate magnesium salt | 0.22mM | Sigma Aldrich | A8960 |
|  | HEPES 1M pH 7.4 | 15mM | Gibco | 15630080 |
|  | FGF-2 basic 145aa | 100ng/mL | Bio-Techne (R&D Systems) | 4114-TC-01M |
|  | TGF-beta | 2ng/mL | Peprotech | AF-100-21C |
|  | 30% human serum albumen (HSA) in water, filter-sterilized (for resuspension of FGF-2) | 0.001% | Sigma | A9080-10ML |
| **mTeSR** | mTeSR™ 1 | N/A | Stemcell Technologies | 85850 |
| **E8** | Essential 8™ Medium | N/A | Gibco | A1517001 |
| **XVIVO**  **differentiation medium** | X-VIVO 15 | 97.8% | SLS (Lonza) | BE02-060F |
|  | GlutaMAX (100x) | 2mM | Gibco | 35050-038 |
|  | 2-Mercaptoethanol (1000x) |  | Gibco | 31350-010 |
|  | IL-3 | 25ng/mL | Invitrogen | PHC0033 |
|  | M-CSF | 50ng/mL | Invitrogen | PHC9501 |
|  | Penicillin-Streptomycin (P/S 100x) | 1% | Gibco | 15140-122 |
| **XVIVO Macrophage medium** | X-VIVO 15 | 97.9% | SLS (Lonza) | BE02-060F |
|  | GlutaMAX (100x) | 2mM | Gibco | 35050-038 |
|  | M-CSF | 50ng/mL | Invitrogen | PHC9501 |
|  | Penicillin-Streptomycin (P/S 100x) | 1% | Gibco | 15140-122 |
| **OXM**  **differentiation medium** | Advanced DMEM/F-12 | 96.3% | Gibco | 12634010 |
|  | GlutaMAX (100x) | 2mM | Gibco | 35050-038 |
|  | HEPES 1M pH 7.4 | 15mM | Gibco | 15630080 |
|  | Human recombinant Insulin solution | 5µg/mL | Sigma | I9278-5ML |
|  | Tropolone | 15µM | Sigma | T89702-1G |
|  | IL-3 | 25ng/mL | Invitrogen | PHC0033 |
|  | M-CSF | 50ng/mL | Invitrogen | PHC9501 |
|  | Penicillin-Streptomycin (P/S 100x) | 1% | Gibco | 15140-122 |
| **OXM Macrophage medium** | Advanced DMEM/F-12 | 96.5% | Gibco | 12634010 |
|  | GlutaMAX (100x) | 2mM | Gibco | 35050-038 |
|  | HEPES 1M pH 7.4 | 15mM | Gibco | 15630080 |
|  | M-CSF | 50ng/mL | Invitrogen | PHC9501 |
|  | Penicillin-Streptomycin (P/S 100x) | 1% | Gibco | 15140-122 |

Complete composition of iPSC and macrophage differentiation medium including supplier and catalogue number. Advanced DMEM/F-12 formulation can be found at the suppliers site at <https://www.thermofisher.com/uk/en/home/technical-resources/media-formulation.227.html> (accessed 11/10/2019)

**Supplementary Table 2: List of metabolites present in both media. Related to Figure 2.**

| Most likely molecule | OXM | | XVIVO | |  | measured | theoretical |
| --- | --- | --- | --- | --- | --- | --- | --- |
|  | Intensity | concentration (mol/L) | Intensity | calculated concentration (mol/L) | Ion | m/z | m/z |
| Tropolone | 8.10E+05 | 1.50E-05 | 1.00E+06 | 1.85E-05 | [M-H] | 121.0293 | 121.0295 |
| Pyruvic acid | 8.00E+08 | 1.00E-03 | 6.70E+06 | 8.38E-06 | [M-H] | 87.009 | 87.0087 |
| Leucine/isoleucine | 3.00E+06 | 4.50E-04 | nd | 0.00E+00 | [M-H] | 130.0524 | 130.0509 |
| L-Aspartic acid | 4.70E+07 | 5.00E-05 | nd | 0.00E+00 | [M-H] | 132.0301 | 132.0302 |
| Glutamic acid | 1.80E+09 | 5.00E-05 | 2.00E+07 | 5.56E-07 | [M-H] | 146.0458 | 146.0458 |
| Thymidine | 7.00E+05 | 1.50E-06 | 1.40E+07 | 3.00E-05 | [M-H] | 241.083 | 241.083 |
| Phenylalanine | 1.80E+05 | 2.15E-04 | 3.00E+04 | 3.58E-05 | [M-H] | 164.0716 | 164.0717 |
| C6 sugars (1) | 2.60E+08 | --- | 5.40E+08 | --- | [M-H] | 179.0559 | 179.05611 |
| C6 sugars (2) | 4.90E+09 | --- | 1.00E+07 | --- | [M-2H] | 89.0243 | 89.024 |
| Tryptophan | 1.10E+05 | 4.40E+05 | 1.60E+07 | 6.40E+07 | [M-H] | 203.0827 | 203.0826 |

Note: the C6 sugars (1) and (2) are different because the intensity ratio (C6 (1)/C6 (2)) is different between OXM and XVIVO.

**Supplemental Experimental Procedures**

**iPSC lines**

The derivation and characterisation of the iPSC lines used in this study is described elsewhere: SFC840-03-03 (Fernandes et al., 2016), SFC841-03-01 (Dafinca et al., 2016) SFC856-03-04 (Haenseler et al., 2017)). All lines were derived from dermal fibroblasts from disease-free donors recruited through StemBANCC (Morrison et al., 2015) and the Oxford Parkinson’s Disease Centre: participants were recruited to this study having given signed informed consent, which included derivation of hiPSC lines from skin biopsies (Ethics Committee: National Health Service, Health Research Authority, NRES Committee South Central, Berkshire, UK, who specifically approved this part of the study (REC 10/H0505/71)). The iPSC lines were all derived using non-integrating Sendai reprogramming vectors (Cytotune, Life Technologies), cultured in mTeSR™1 on hESC-qualified Matrigel-coated plates (BD, #356234), passaging as clumps using 0.5 mM EDTA in PBS (Beers et al., 2012). Large-scale, low-passage frozen SNP-QCed batches were used for experiments to ensure consistency.

**Cell culture reagent preparation: OXE8**

Ascorbic acid was dissolved in PBS to make a 0.22 M stock and aliquoted at 1mL for storage at -20^o^C indefinitely, or 4^o^C for up to 3 months. FGF-2 was reconstituted according to manufacturer’s instructions and a 1 mg/mL stock made in 0.1% HSA and aliquoted at 50 µL for storage at -80^o^C. 10 µg Lyophilised TGF-β was first centrifuged to pellet the powder before reconstitution in 100 µL ultra-pure sterile water. This was then transferred into 4.8 mL 0.1% HSA solution to make a 2 µg/mL stock and aliquoted at 0.5 mL for storage at -80^o^C. Both FGF-2 and TGF-β once thawed are considered stable for 2 weeks at 4^o^C. Avoid freeze/thaw cycles. To make OXE8 was as follows: 500 mL aDMEM/F-12, 5 mL GlutaMAX, 25 µL Heparin, 0.5 mL Ascorbic acid, 7.5 mL HEPES pH 7.4, 50 µL FGF-2, and 0.5 mL TGF-β.

**Cell culture reagent preparation: OXM**

The desired mass of powdered Tropolone was dissolved in ultra-pure water and filter sterilised by passing through a 0.22 µm filter. It was aliquoted at 1.5 mL and stored at -20^o^C indefinitely, or 4^o^C for up to 3 months. IL-3 was reconstituted according to manufacturer’s instructions in ultra-pure sterile water and aliquoted at 0.5 mL for storage at -80^o^C. Once thawed, it is considered stable for 1 month at 4^o^C. M-CSF was reconstituted as needed according to manufacturer’s instructions in 20 mM Tris-HCL pH 8.0. Each vial was resuspended in 1 mL and could be stored at for 1 month at 4^o^C or stored at -80^o^C. To make OXM differentiation media was as follows: 500 mL aDMEM/F-12, 5 mL GlutaMAX, 5 mL P/S, 0.25 mL Insulin solution, 7.5 mL HEPES pH 7.4, 0.75mL Tropolone, 0.5 mL M-CSF, and 0.25 mL IL-3. To make OXM macrophage medium Tropolone and IL-3 were excluded.

**Cell culture**

iPSC were cultured in their stated media (Supplementary Table 2) on Geltrex™ (Gibco, #A1413201)-coated tissue culture dishes and passaged using TrypLE™ Express (Gibco, #12604013). For 24 hours after plating, media was supplemented with 10 µM Y-27632 (Abcam, ab120129). Vero-76 and HEK293T cells were cultured in DMEM made to 10% FBS (Sigma, #F9665) and 1% Pen/Strep (Gibco, #15140122). All cells were incubated at 37^o^C, 5% CO_2_.

**Macrophage differentiation**

iPSC were differentiated into macrophages using a protocol based on that previously described (van Wilgenburg et al., 2013). The updated method is as follows. Aggrewell™ 800 plates (Stemcell technologies, #34815) were prepared by addition of 0.5 mL Anti-Adherence Rinsing solution (Stemcell technologies, #07010) and centrifugation at 3000g for 3 minutes to remove bubbles from the microwells. Rinse solution was then aspirated and replaced with 1 mL of 2X concentrated EB medium (1X EB media: OXE8 media supplemented with 50 ng/mL BMP4 (Peprotech, #PHC9534), 50 ng/mL VEGF (Peprotech, #PHC9394), and 20 ng/mL SCF (Miltenyi Biotec, #130-096-695) supplemented with 10 µM Y-27632. iPSC were resuspended by washing with PBS, incubating in TrypLE Express for 3-5 minutes at 37^o^C, 5% CO_2_, followed by gentle lifting in Advanced-DMEM/F12 to achieve single cell suspension. Cells were counted and pelleted by centrifugation at 400g for 5 minutes. After centrifugation, cells were resuspended at 4x10^6^ cells/mL in OXE8 supplemented with 10 µM Y-27632 and 1 mL added to the Aggrewell. The Aggrewell plate was then spun at 100g for 3 minutes with no braking to encourage even distribution of cells across microwells. Cells were incubated for 4 days at 37^o^C, 5% CO_2_ with daily feeding of 75% media change with EB medium, by aspiration of 1 mL by pipette and gentle addition of 1 mL fresh media twice to avoid disturbance to the microwells. After 4 days, EBs were lifted from the plate using a Pasteur pipette and passed over a 40 µm cell strainer to remove dead cells, before washing into tissue culture plate with differentiation media (Supplementary Table 2). EBs were divided evenly into 2 T175 flasks and topped up to 20 mL with differentiation medium. Differentiation cultures were incubated at 37^o^C, 5% CO_2_, with weekly feeding of an additional 10 mL until macrophage precursors (PreMac) cells started to be produced. After this point PreMac cells were collected weekly and a minimum of equal volumes of media to the volume removed were replaced into the differentiation cultures. Each harvest involved a 25-50% media change. Harvested cells were either used directly or plated in appropriately sized tissue culture plates for further culturing in XVIVO or OXM Macrophage medium (Supplementary Table 2) for a further 7 days, with a 50% media change on day 4.

**Cell count, size, and viability measurements**

Day 7 macrophages were resuspended by replacing media with StemPro™ Accutase™ Cell Dissociation Reagent (Stemcell technologies, #A1110501) and incubating 3-5 minutes at 37^o^C, 5% CO_2_. Cells lifted in Accutase were diluted in macrophage media 1:10 to maintain accurate viability throughout measurements. Cell count, size, and viability were measured using the Nucleocounter® NC-3000 (Chemometec) after staining for live/dead cells using Solution-13 AO-DAPI stain (Chemometec, #910-3013) mixed 1:20 dye to cell suspension.

**Flow Cytometry**

Cells were lifted using Accutase as described above as no difference in surface marker levels using this method versus 5 mM EDTA/12 mM Lidocaine has been observed (Carter et al 2009, Haenseler et al 2018). Cells in suspension were kept at 4^o^C or on ice throughout staining. For surface marker expression, cells were stained directly without fixation in FACS buffer (PBS supplemented to 1% FBS, 10 µg/mL human-IgG (Sigma, #I8640-100MG), and 0.01% Sodium azide). For total marker staining, cells were first fixed for 10 minutes in 2% PFA over ice. They were then permeabilised by treatment with 0.1% saponin in PBS for 1 hour at room temperature. Staining was then carried out in FACS buffer supplemented with 0.1% saponin. Isotype controls with the same fluorophores from the same company were used at the same concentration. Fluorescence was measured using the BD LSRFortessa™ X-20 (BD Biosciences) and analysed using FlowJo version 10. Antibodies used for flow cytometry of surface markers were all mouse primary conjugated antibodies. Antibodies used were as follows: CD4 (APC, clone 11830, R&D systems, #FAB3791A), CD11b (APC, clone ICRF44, Bio-legend, #301310) CD206 (APC, clone 15-2, Bio-legend, #321109), CCR5 (PE, clone 45531, R&D, #FAB182P), CD16 (PE, clone PPV-06, Immunotools, #21279164), CD86 (PE, clone IT2.2, Bio-legend, 305405), CD163 (PE, clone 215927, R&D, #FAB1607P), CXCR4 (PE, clone 44717, R&D, FAB173P), CD14 (FITC, clone MEM-18, Immunotools, #21275514), CD45 (FITC, clone MEM-28, Immunotools, #21270453X2), CD68 (FITC, clone Ki-M7, BIO-RAD, #MCA2375F), CD86 (FITC, clone BU63, Immunotools, #21480863), HLA-DR (FITC, clone MEM-12, Immunotools, #21388993), IgG1κ (APC, clone MOPC-21, Bio-legend, #400120), IgG2a (APC, clone 20102, R&D, #IC003A), IgG1 (PE, clone PPV-06, Immunotools, #21275514), IgG2bk (PE, clone MPC-11, Bio-legend, #400311), IgG1 (PE, clone 11711, R&D, #IC002P), IgG2b (PE, clone 133303, R&D, #IC0041P), IgG1 (FITC, clone PPV-06, Immunotools, #21335013), and IgG1 (FITC, BIO-RAD, #MCA928F). Staining for KI67 used a mouse primary antibody (Abcam, #ab15580) or isotype control primary IgG (Abcam, #ab172730) followed with a goat anti-mouse Alexa-488 conjugated secondary antibody (Invitrogen, #A-21131).

**Cytokine and Chemokine release**

As previously described (van Wilgenburg et al., 2013), 2.8x10^5^ macrophages plated in a 24-well plate were stimulated with 100 ng/mL LPS (Invivogen, #tlrl-eklps) and 20 ng/mL IFNγ (Gibco, #PHC4031 ), or 50 ng/mL IL-4 (Gibco, #PHC0044) for 16 hours after which supernatants were collected and spun at 400g for 5 minutes to remove dead and floating cells, and stored at -80C until required. Cytokine release was measured using the Proteome Profiler Human Cytokine Array Kit (R&D systems, #ARY005B) modified to use IRDye 800CW Streptavidin (LI-COR, #926-32230) according to R&D systems protocol (<https://www.rndsystems.com/resources/technical/use-proteome-profiler-arrays-li-cor-detection>) for visualizing results on the LI-COR Odyssey 9260 and quantification using Image Studio Lite version 5.2.

**Phagocytosis**

Macrophages were cultured at 2x10^5^ cells/well in a 24-well plate. Control cells were pre-treated for 1 hour with 10 µM Cytochalasin D (Cambridge Bioscience, #11330-1 mg-CAY). 12.5 µg Zymosan A (*S. cerevisiae*) BioParticles™ Alexa Fluor-488 (ThermoFisher, #Z23373) were added to each well of cells and incubated for 30 minutes at 37^o^C, 5% CO_2_. Particles not taken up were quenched with 0.025% (v/v) Trypan Blue (Sigma, #T8154) in PBS before lifting the cells with Accutase as described above. Cells were fixed with 4% paraformaldehyde (PFA) (Alfa Aesar, #J61899.AK) before measuring fluorescence with the BD LSRFortessa™ X-20.

**Lentivirus preparation and Infection assay**

Lentiviral vectors were produced using PEI-mediated transfection of HEK 293T cells with pNL4.3R-E-eGFPT2ANef and pBa-L. Lentiviral containing supernatant was harvested at 48 hours, filtered, and concentrated 100X using PEG6000.

5x10^4^ macrophages plated in 96-well format were infected with the stated volumes of virus diluted to 100 µL and incubated for 72 hours. Cells were resuspended by pipetting after using 5 mM EDTA in PBS at 4^o^C for 30 minutes, and fixed in 1% PFA. GFP positive cells were measured by flow cytometry on the BD LSRFortessa™ X-20.

**One-step growth of Zika virus (ZIKV) in macrophages**.

ZIKV isolate MR1766, Uganda 1947, was obtained from EVAg, the European Virus Archive. 2x10^5^ macrophages in a 24-well dish were infected with 2 x 10^6^ pfu of ZIKV. After 4 hours at 37^o^C, 5% CO_2_, supernatants were aspirated and replaced with 700 µL fresh media, and incubation continued. 200 µL samples of supernatant were harvested at intervals from 6h to 48 hours post-infection, clarified by centrifugation and stored at -80^o^C.

**Titration of ZIKV infectivity**.

Virus-containing supernatants were serially diluted 10-fold and 100 µL added to each well of a 24-well plate. 2.5x10^5^ Vero-76 Cells in maintenance media (DMEM containing 1% FBS and 1% Pen-Strep to maintain cell viability without proliferation) were then added. Cells were incubated with virus for 2 hours at 37^o^C, 5% CO_2_, before adding a semi-solid overlay of 1.5% Carboxymethyl cellulose sodium salt, low viscosity (Sigma, #C5678) in Vero maintenance media and returned to 37^o^C, 5% CO_2_. At 72 hours, the semi-solid overlay was removed, the cell monolayers washed with PBS, and plaques were revealed with Amido black stain.

**RNA-seq sample preparation, library construction, and data generation**

RNA was extracted from 1x10^6^ macrophages (SFC840-03-03) differentiated from PreMac cells harvested at week 8, 10, and 12 of differentiation, using the RNeasy Mini Kit (QIAGEN, # 74104) according to manufacturer’s directions, including optional DNase treatment step. Samples were sent to Novogene on dry ice for analysis. RNA degradation and contamination was monitored on 1% agarose gels. RNA purity was checked using the NanoPhotometer® spectrophotometer (IMPLEN, CA, USA). RNA integrity and quantitation were assessed using the RNA Nano 6000 Assay Kit of the Bioanalyzer 2100 system (Agilent Technologies, CA, USA). A total amount of 1 µg RNA per sample was used as input material for the RNA sample preparations. Sequencing libraries were generated using NEBNext® Ultra TM RNA Library Prep Kit for Illumina® (NEB, USA) following manufacturer’s recommendations and index codes were added to attribute sequences to each sample. Briefly, mRNA was purified from total RNA using poly-T oligo-attached magnetic beads. Fragmentation was carried out using divalent cations under elevated temperature in NEBNext First Strand Synthesis Reaction Buffer (5X). First strand cDNA was synthesized using random hexamer primer and M-MuLV Reverse Transcriptase (RNase H-). Second strand cDNA synthesis was subsequently performed using DNA Polymerase I and RNase H. Remaining overhangs were converted into blunt ends via exonuclease/polymerase activities. After adenylation of 3’ ends of DNA fragments, NEBNext Adaptor with hairpin loop structure were ligated to prepare for hybridization. In order to select cDNA fragments of preferentially 150~200 bp in length, the library fragments were purified with AMPure XP system (Beckman Coulter, Beverly, USA). Then 3 µl USER Enzyme (NEB, USA) was used with size-selected, adaptorligated cDNA at 37 °C for 15 min followed by 5 min at 95 °C before PCR. Then PCR was performed with Phusion High-Fidelity DNA polymerase, Universal PCR primers and Index (X) Primer. At last, PCR products were purified (AMPure XP system) and library quality was assessed on the Agilent Bioanalyzer 2100 system. The clustering of the index-coded samples was performed on a cBot Cluster Generation System using PE Cluster Kit cBot-HS (Illumina) according to the manufacturer’s instructions. After cluster generation, the library preparations were sequenced on an Illumina platform and paired-end reads were generated.

**RNA-Seq data analysis**

Raw reads were pre-processed using fastp (v.0.20.1) software (Chen et al., 2018). In addition to the default filtering parameters, polyN sequences in read tails and adapter sequences were trimmed, and mismatched base pairs were corrected by overlap analysis. Filtered and trimmed reads were mapped to a full decoy-aware transcriptome using the Gencode version 34 reference human transcriptome and GRCh38 primary assembly genome. Salmon (v.1.2.1) selective alignment was used for transcript quantification with optional flags correcting for GC and random hexamer priming biases (Srivastava et al., 2019). All downstream analyses were done using the R (v.4.0) programming language (R Core Team 2020). Transcript abundance estimates in TPMs were summarised to the gene level using the tximport (v.1.16.0) package to correct for sample specific transcript length biases (Soneson et al., 2016). Lowly expressed genes with length-scaled abundance estimates less than 10 in 3 samples were filtered out. Differential expression was tested using limma-voom (Law et al., 2014). GO enrichment analysis was conducted using the EGSEA package (Alhamdoosh et al., 2017). GO terms with less than 10 genes were excluded from the enrichment analysis. The background gene set used for the analysis was the total number of the unique genes observed in the experiment. A publicly available dataset of human *ex vivo* and iPSC-derived microglia was processed as above and used as a reference (Abud et al., 2017). Principal component analysis was performed with the prcomp package to visualise the level of transcriptomic similarity amongst the samples from both studies.

**Liquid chromatography-mass spectrometry (LC-MS) analysis of media**

Samples of the media were prepared for LC-MS analysis as described before (Ebrahimi et al., 2020). Briefly, 500 µL of each media was filtered using Amicon Ultracentrifugal filters (3 or 10 kDa cut off). The flow through was used for analysis of metabolites using high-resolution negative ion LC-MS (accuracy better than 5 ppm). The setting and methods were as described previously (Alldritt et al., 2019; Ebrahimi et al., 2020). The total ion chromatograms were analysed using MesReNova software. The MS peaks were putatively assigned using and the human metabolome database (HMDB). The concentrations in the XVIVO were estimated based on the known concentration of different metabolites in the OXM media the ratio of the intensities for each molecule (Concentration in XVIVO = Concentration in OXM*(intensity XVIVO / intensity OXM).

**Quantification of glucose concentration**

A modification of the protocol for the detection of glucose using the Glucose (HK) Assay Kit (Supelco, # GAHK20-1KT) from culture media was done as follows: 100 µL of culture media were diluted 1:10 and 1:20 in dH2O. 20 µL of each dilution were added onto a well of a 96-well flat-bottom cell culture treated plate. Glucose solution was freshly made up by resuspending the vial using 20 mL of dH2O. 100 µL of glucose solution were added per well. Plates were incubated at 35^o^C for 15 minutes before measuring absorbance at 340 nm using a plate reader. Values were compared to known values from a standard curve. The standard curve was done using seven serials dilutions (top 2.5 mM) of media culture with known composition (Dulbecco's Modified Eagle Medium – 25 mM). Each sample was run in duplicates on the plate.
