## Supplementary figures and images for "Differentiation of Human induced Pluripotent Stem Cells to Authentic Macrophages using Fully Defined, Serum Free, Open Source Media"

### Supplementary Figures

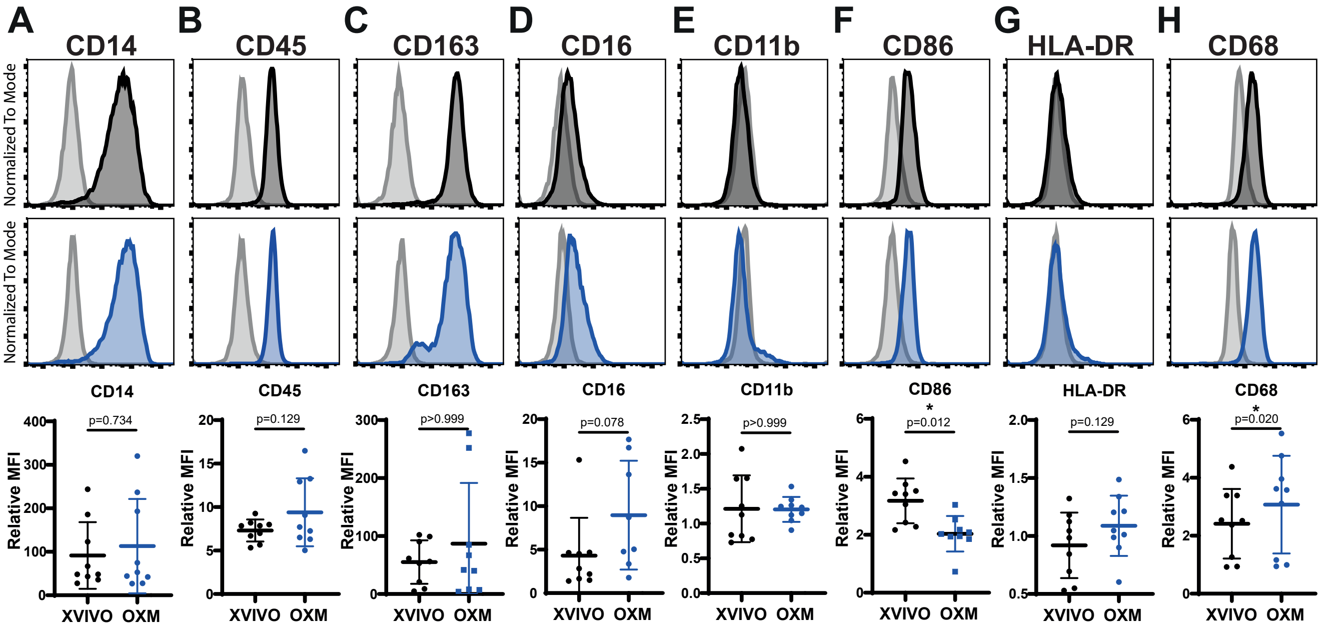

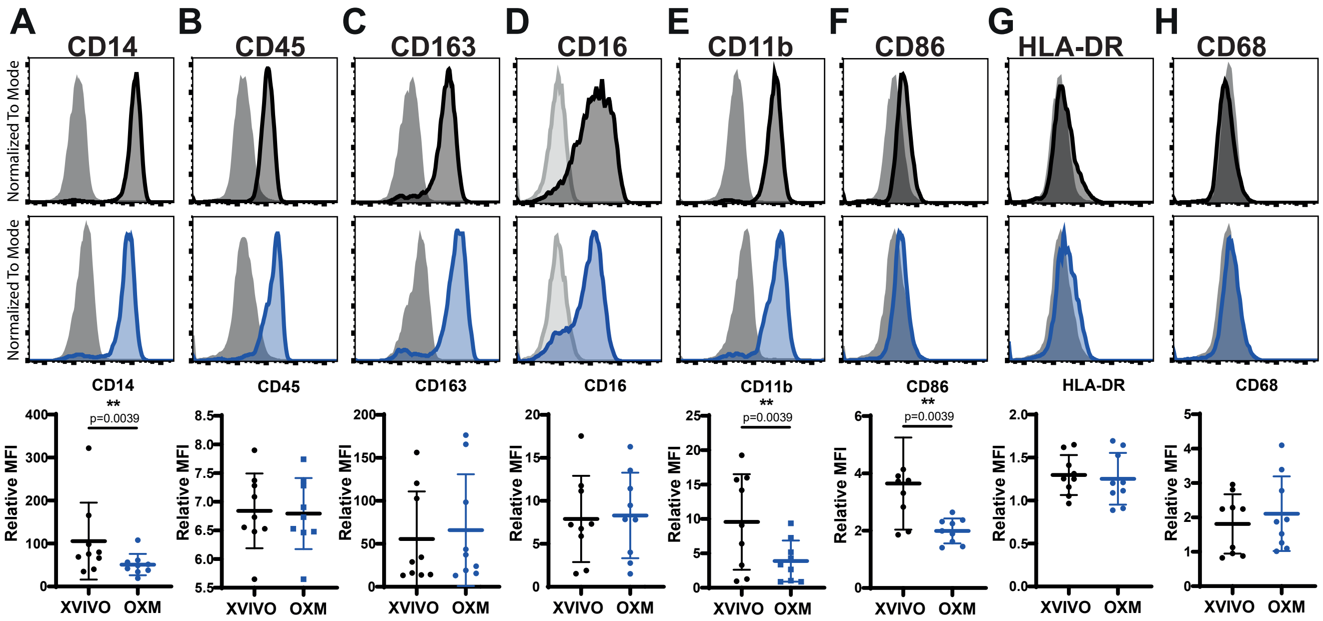

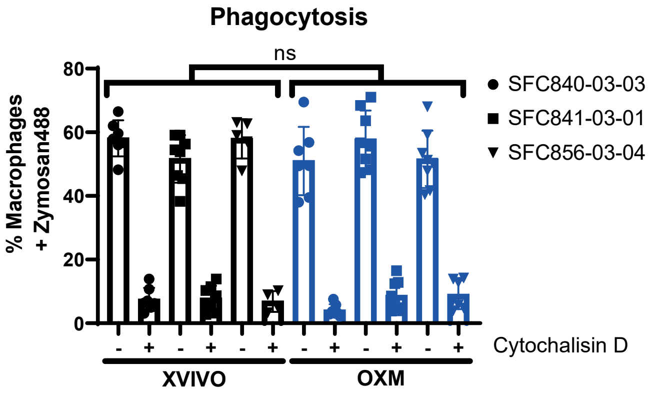

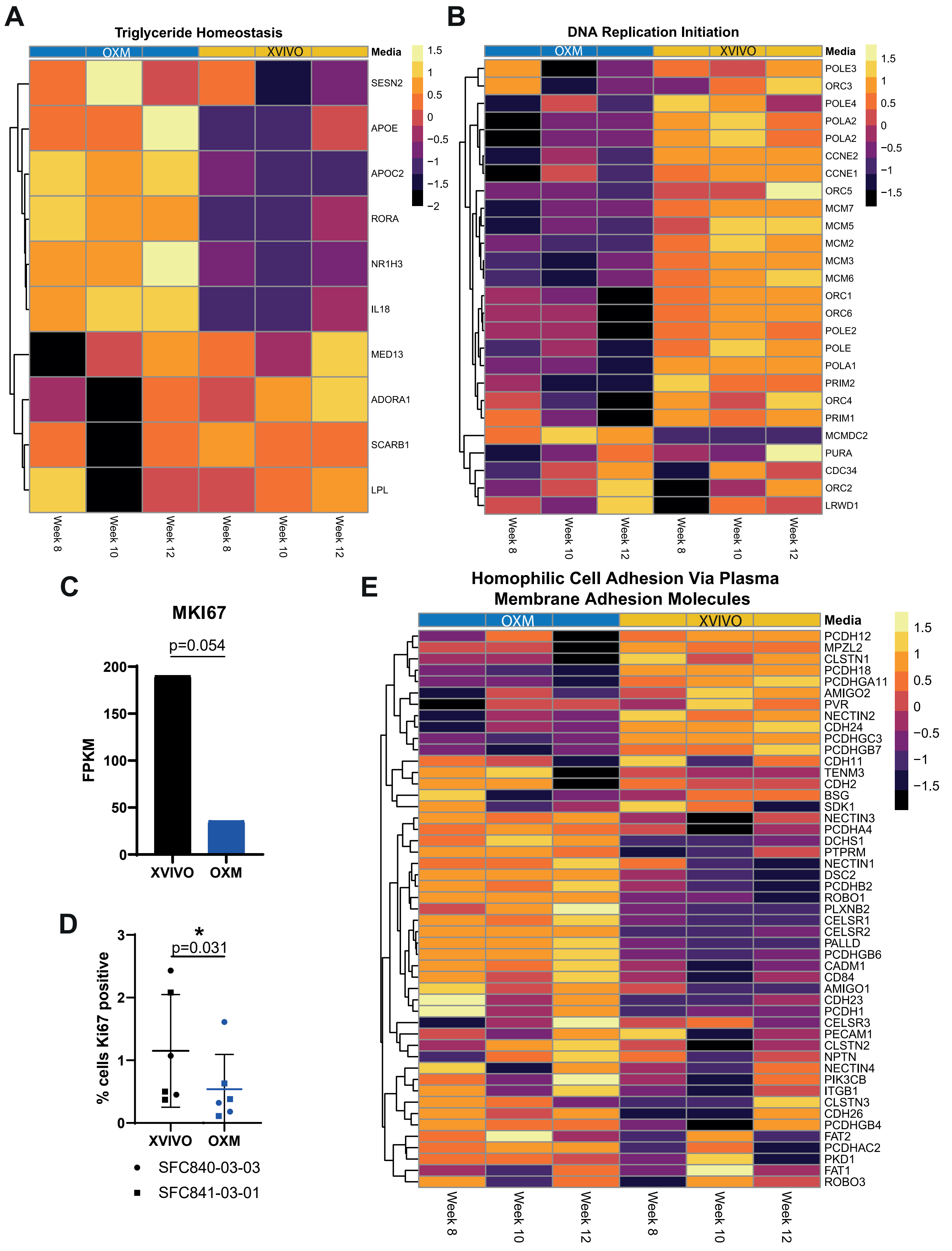
